# Chronic postnatal chemogenetic activation of forebrain excitatory neurons modulates adult glial function and metabolism in male mice

**DOI:** 10.64898/2026.05.16.725656

**Authors:** Amartya Pradhan, Sthitapranjya Pati, Kamal Saba, Pratik R Chaudhari, Praachi Tiwari, Darshana Kapri, Angarika Balakrishnan, Anant B Patel, Vidita A Vaidya

## Abstract

Early adversity increases vulnerability for adult psychopathology. Across multiple pre-clinical models of early adversity, there are reports of glial dysfunction and disrupted amino acid neurotransmission, along with maladaptive behavioral responses in adulthood. Disrupted G-protein coupled receptor signaling is known to phenocopy specific consequences of early life adversity. Enhanced Gq signaling in the forebrain excitatory neurons in early postnatal life programs anxio-depressive behaviors in adulthood, accompanied by altered neuronal glutamate and GABA metabolism in mouse models. We hypothesized that enhancing Gq signaling in forebrain excitatory neurons in early postnatal life may also impact glial function in adulthood. Our results show that postnatal hM3Dq-mediated chemogenetic activation of CaMKIIα-positive forebrain excitatory neurons not only increases anxiety-like behavior, but also evokes bidirectional transcriptional regulation of multiple glia-associated genes in the neocortex and hippocampi. While *Gfap, Aldh1l1, S100*β, *Eaat1, Eaat2* and *Eaat3*, mRNA levels were reduced in the neocortex, they were enhanced in the hippocampus, and a similar pattern was noted for GFAP protein levels. Transient, postnatal chemogenetic activation of CaMKIIα-positive neurons did not alter astrocyte cell density in both the neocortex and the hippocampus. Using (^1^H-(^13^C)) NMR spectroscopy, we observed a significant decline in astrocyte-specific glutamate and GABA neurotransmitter turnover, and a reduction in astrocyte metabolic flux within the neocortex and the hippocampus in adulthood in animals with a history of postnatal chemogenetic activation of forebrain excitatory neurons. Our findings indicate that chemogenetically driving Gq signaling transiently during the postnatal window in forebrain excitatory neurons results in persistent changes well into adulthood, with enhanced anxiety-like behaviors and disrupted glial function and metabolism, phenocopying specific changes in glial function noted following early adversity.

## INTRODUCTION

Evidence from clinical and pre-clinical studies indicates that early adversity is a common risk factor for the development of several neuropsychiatric disorders, ranging from anxiety, depression, and schizophrenia to substance abuse disorders (Felitti *et al*., 1998; Ansorge *et al*., 2007; Bale *et al*., 2010; Targum & Nemeroff, 2019). Amongst the mechanisms implicated in programming the persistent behavioral consequences of early adversity, is perturbed monoaminergic neurotransmission and G protein-coupled receptor (GPCR) signaling during key early epochs of life. Preclinical studies suggest that aberrant GPCR signaling, especially that downstream of Gq-coupled and Gi-coupled monoaminergic GPCRs, in particular, the Gq-coupled serotonin_2A_ (5-HT_2A_) (Weisstaub *et al*., 2006; Sarkar *et al*., 2014) and the Gi-coupled serotonin_1A_ (5-HT_1A_) receptor (Gross *et al*., 2002; Richardson-Jones *et al*., 2010, 2011; Garcia-Garcia *et al*., 2014), contributes to the programming of altered trait anxiety in adulthood. Notably, the Gq-coupled 5-HT_2A_ receptor exhibits altered signaling in diverse models of early adversity, including maternal immune activation (Holloway *et al*., 2013; Malkova *et al*., 2014), maternal separation (Benekareddy *et al*., 2010, 2011; Sood *et al*., 2018), and postnatal fluoxetine administration (Sarkar *et al*., 2014; Ghai *et al*., 2026). Prior studies indicate that blockade and regulation of Gq-coupled GPCR signaling within distinct circuits has differential effects on anxio-depressive behaviors. For instance, while transient chemogenetic activation of Gq signaling in CaMKIIα-positive forebrain excitatory neurons during the postnatal window can enhance anxio-depressive behaviors (Pati *et al*., 2020), driving Gq-signaling in the same window, albeit with a pan-neuronal marker (hSyn) restricted to the prefrontal cortex, decreased anxio-depressive behaviors associated with natural and pharmacological models of early life adversity (Soiza-Reilly *et al*., 2018; Teissier *et al*., 2019). Postnatal hM3Dq-mediated chemogenetic activation of all PV-positive neurons evoked a reduction in anxio-depressive behaviors in adulthood, in a task- and sex-dependent manner (Banerjee *et al*., 2022). These studies assessing the long-term impact of transient postnatal activation of Gq signaling have predominantly focused on the impact on adult mood-related behavioral tasks, and associated molecular, cellular, and metabolic consequences in neuronal populations. There remains a paucity of information on whether such perturbations of Gq-signaling in neurons can evoke any long-lasting impact within the milieu, particularly, on the other major cell type in the brain, astrocytes.

Astrocytes perform critical and diverse functions in the brain, including neurotransmitter cycling, maintaining the blood-brain barrier, and regulating synaptic plasticity. Accumulating evidence suggests that glial dysfunction, in particular astrocyte structural and functional pathology, is implicated in contributing to the profound alterations in adult mood-related behaviors associated with early adversity (Abbink *et al*., 2019; Teissier *et al*., 2019; Bansal *et al*., 2024; Guayasamin *et al*., 2025). Preclinical studies have focused on structural alterations of glial cells and glial cell-associated gene expression in the context of early adversity; however, the functional consequences on non-neuronal cells, including astrocytes, remain poorly understood. To address these gaps, we utilized a chemogenetic strategy with a CaMKIIα-tTA::TetO-hM3Dq bigenic mouse line, where the excitatory Designer Receptor Exclusively Activated by Designer Drug (DREADD), hM3Dq, was selectively expressed in forebrain CaMKIIα-positive excitatory neurons. DREADD-specific Gq signaling was transiently enhanced in the critical early postnatal window (P2 to P14) with the administration of the synthetic DREADD ligand, clozapine N-oxide (CNO). We hypothesized that transiently enhancing Gq signaling in forebrain excitatory neurons during the postnatal window may exert persistent impacts on glial function and metabolism in adulthood, especially in astrocytes. We find that our chemogenetic perturbation enhanced anxiogenic behaviors, recapitulating findings from our previous study (Pati *et al*., 2020). Concomitantly, we observed bidirectional, brain region-specific changes in glia and glia-associated gene and protein expression, without evidence for any alterations in astrocyte cell numbers in the neocortex and hippocampi. Using (^1^H-(^13^C)) nuclear magnetic resonance (NMR) spectroscopy, we demonstrate that transient postnatal increases in Gq signaling in forebrain excitatory neurons program persistent alterations in glutamate and GABA turnover in astrocytes, along with a reduction in metabolic rate. Our findings indicate that Gq signaling perturbation in excitatory neurons during postnatal life can program long-lasting changes in glial function and metabolism in key limbic brain regions.

## MATERIALS AND METHODS

### Animals

All animals were bred and raised in the Tata Institute of Fundamental Research (TIFR), Mumbai, India, animal house facility on a 12 hr light/dark cycle (7 AM to 7 PM) with *ad libitum* access to food and water. Wildtype C57BL/6J mice or bigenic CaMKIIα-tTA::tetO-hM3Dq were used for all experiments. Bigenic CaMKIIα-tTA::tetO-hM3Dq mice were generated by mating CaMKIIα-tTA mice (gifted by Dr. Christopher Pittenger, Department of Psychiatry, Yale School of Medicine, New Haven, USA) (Mayford *et al*., 1996) with tetO-hM3Dq mice (Tg(tetO-CHRM3*)1Blr/J; Strain No. 014093, The Jackson Laboratory, USA), and genotypes were confirmed with PCR-based analysis. Primiparous dams were kept as dyads prior to single-housing one/two days before parturition and were provided with paper shredding as nesting material. Litter size was restricted to 6-8 pups per litter.

^1^H-[^13^C]-NMR spectroscopy experiments from tissue samples were carried out at the Centre for Cellular and Molecular Biology (CCMB), Hyderabad, India. Animal handling and experiments were carried out as per the guidelines of the Committee for Control and Supervision of Experiments on Animals (CCSEA), Government of India, and were approved by the TIFR (IAEC/2019-4) and CCMB (IAEC-32/2018) animal ethics committees. Care was taken across all experiments to minimize animal suffering and restrict the number of animals used.

### Drug administration

Litters were randomly assigned to either postnatal CNO (PNCNO) treatment or vehicle treatment groups. To assess the role of enhanced DREADD-mediated Gq signaling within forebrain excitatory neurons during the postnatal window, CaMKIIα-tTA::tetO-hM3Dq mouse pups were orally administered the DREADD-agonist, Clozapine N-Oxide (CNO; Catalog No. 4936, Tocris Bioscience, UK) at a dose of 1mg/kg or vehicle (5% aqueous sucrose solution) once a day through a micropipette (0.5-10 *µ*L; Catalog No. 3123000020, Eppendorf, Germany) from postnatal day 2 (P2) to P14. To minimize any handling-related stress, the oral administration was completed within 3 minutes to avoid prolonged separation from the dam. The postnatal treatment window (P2-P14) was kept consistent with our prior study on the postnatal chemogenetic activation of CaMKIIα-positive forebrain excitatory neurons (Pati *et al*., 2020). The weight profile of pups was monitored throughout the postnatal drug administration paradigm. After weaning (P24-27), male and female mice were separated and group-housed (3-4 animals per cage) and left undisturbed until adulthood (4-6 months), except for routine mouse colony maintenance. Adult male bigenic CaMKIIα-tTA::tetO-hM3Dq mice were used for behavioral, molecular, cytological, and metabolic analyses.

### Behavioral analysis

Bigenic vehicle- and CNO-treated CaMKIIα-tTA::tetO-hM3Dq adult male mice were subjected to behavioral assays to assess anxiety-like behaviors on the elevated plus maze (EPM) test. All trials were recorded for 10 minutes using an overhead camera at 25 frames per second (fps), and behavioral trajectories were tracked using the automated behavioral tracking and analysis software, EthoVision XT 11 (Noldus, the Netherlands). The EPM test was carried out on a plus-shaped platform consisting of two open and closed arms (30 cm x 5 cm each) elevated 50 cm above the ground. The walls of the closed arms of EPM were 15 cm high. The animals were introduced in the center of the maze, facing either of the open arms at random. The arena was thoroughly cleaned before the onset of every trial. To assess anxiety-like behavior on the EPM test, the total distance travelled in the maze, percent distance travelled in open arms, percent time spent in open and closed arms, as well as the number of entries to open and closed arms, were measured for each trial.

### Western blotting

To assess the expression of HA-tagged hM3Dq-DREADD and to determine the effect of chronic postnatal chemogenetic activation of CaMKIIα-positive neurons on protein expression of glial markers within the forebrain, western blotting assays were performed on neocortical and hippocampal tissues derived from vehicle- and CNO-treated bigenic CaMKIIα-tTA::tetO-hM3Dq adult male mice. Animals were anesthetized by CO_2_ inhalation and killed by rapid decapitation. Neocortical and hippocampal tissues were microdissected in ice-cold phosphate buffer saline (PBS) solution, snap-frozen with liquid nitrogen, and stored at -80°C. Frozen tissues were homogenized in radioimmunoprecipitation assay (RIPA) buffer (containing 10 mM Tris-Cl (pH 8.0), 1 mM EDTA, 0.5 mM EGTA, 1% Triton X-100, 0.1% sodium deoxycholate, 0.1% SDS, 140 mM NaCl, and protease and phosphatase inhibitors) using a Dounce homogenizer. Protein concentration was estimated using the QuantiPro BCA Assay Kit (Catalog No. QPBCA, Sigma-Aldrich, USA). The lysates (30-50 μg) were resolved on a 10% sodium dodecyl sulphate (SDS) polyacrylamide gel before transferring them onto polyvinylidene fluoride (PVDF) membranes (Merck Millipore, Germany). PVDF blots were subjected to a blocking step in either 5% non-fat dry milk or 5% bovine serum albumin prepared in Tris-buffered saline containing Tween 20 (TBST) for 1 hr, and then incubated overnight with either of the following primary antibodies: rabbit anti-HA (1:1000; Catalog No. H6908, Sigma-Aldrich, USA), rabbit anti-GFAP (1:2000; Catalog No. G9269, Sigma-Aldrich, USA), or rabbit anti-β-actin (1:12,000; Catalog No. AC026, ABclonal Technology, USA) antibody. Following subsequent washes with TBST, blots were incubated with HRP-conjugated goat anti-rabbit secondary antibody (1:6000; Catalog No. AS014, ABclonal Technology, USA) for 1 hr. Following washes, the signal was visualized on an Amersham Imager 800 (GE Life Sciences, USA) using a western blotting detection kit (WesternBright ECL, Catalog No. K-12045, Advansta Inc., USA). The densitometric quantitative analysis was performed using ImageJ software.

### Quantitative polymerase chain reaction

To understand the influence of chronic postnatal hM3Dq DREADD-mediated activation of CaMKIIα-positive forebrain excitatory neurons on gene expression of glial markers and glia-associated genes within the forebrain, quantitative polymerase chain reaction (qPCR) assays were performed on the neocortex and hippocampi derived from vehicle- and CNO-treated bigenic CaMKIIα-tTA::tetO-hM3Dq adult male mice. Animals were anesthetized by CO_2_ inhalation and killed by rapid decapitation. Neocortical and hippocampal tissues were microdissected in ice-cold phosphate buffer saline (PBS) solution, snap-frozen with liquid nitrogen, and stored at -80°C. RNA was extracted using the TRIzol reagent (Catalog No. 15596026, Invitrogen, USA), quantified using a NanoDrop spectrophotometer (NanoDrop One; Catalog No. ND-ONE-W, ThermoFisher Scientific, USA), and further subjected to reverse transcription reaction to generate cDNA using the PrimeScript RT Reagent Kit (Catalog No. RR037A, Takara Bio, Japan). The master mix for qPCR was prepared using the synthesized cDNA, primers specific to genes of interest **(Table 1)**, and SYBR Green kit: KAPA SYBR FAST qPCR Master Mix (Catalog No. KK4601, KAPA Biosystems, Switzerland) or SsoAdvanced Universal SYBR Green Supermix (Catalog No. 1725271, Bio-Rad, USA). The qPCR was performed using a real-time PCR detection system (CFX96; Bio-Rad, USA). Primers were designed using the NCBI Primer-BLAST tool (National Center for Biotechnology Information, USA). The C_t_ value for a particular gene was normalized to the 18S rRNA housekeeping gene. The qPCR data were analyzed using the ^ΔΔ^C_t_ method as previously described (Bookout *et al*., 2003).

**Table 1:**
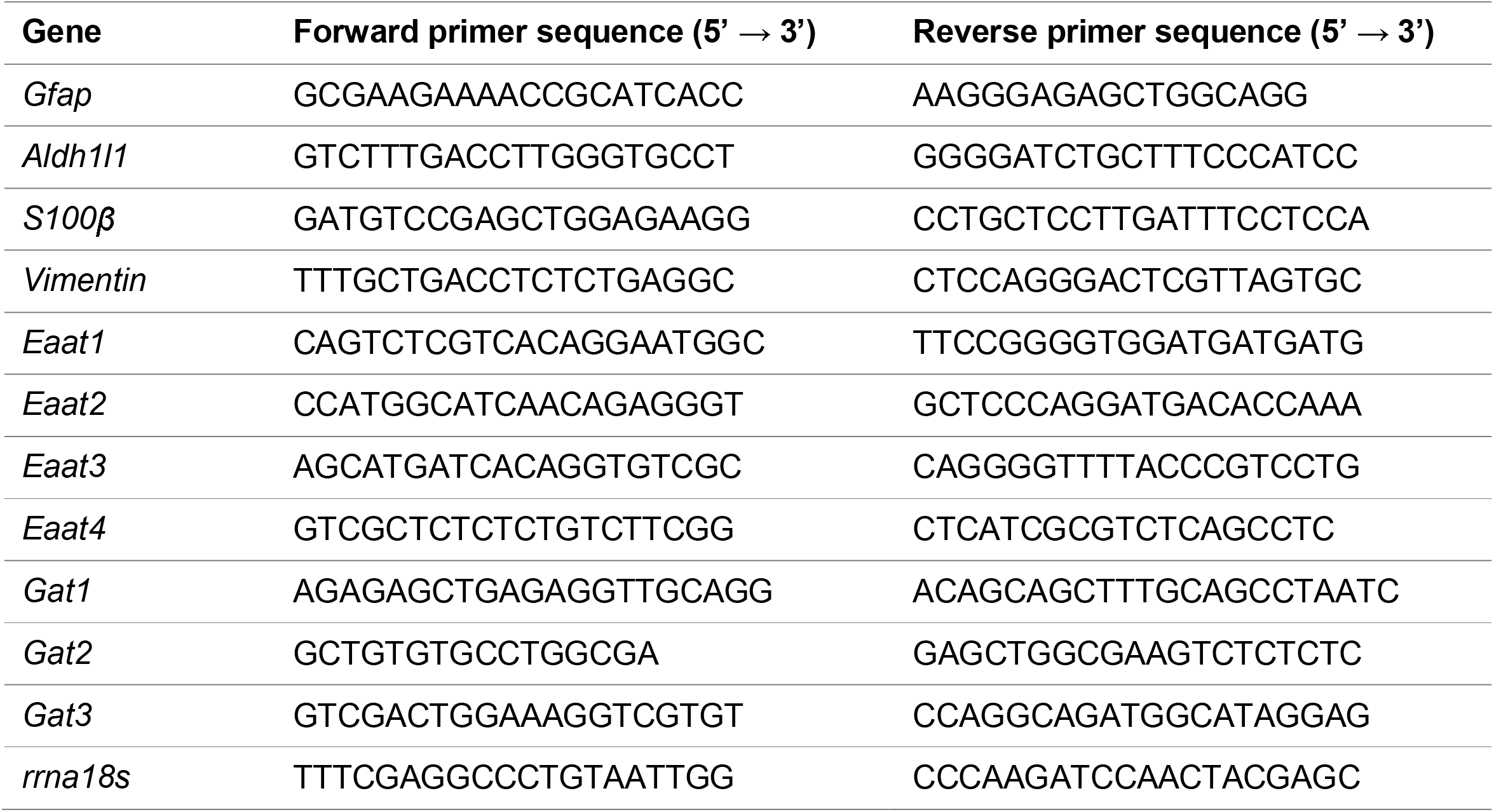
List of qPCR oligonucleotide primer sequences.

### Immunofluorescence

Brains were harvested from vehicle- and CNO-treated bigenic CaMKIIα-tTA::tetO-hM3Dq adult male mice by transcardial perfusion using 4% paraformaldehyde. Serial coronal brain sections (50 *µ*m) were obtained using a vibratome (Leica VT1000 S; Leica Biosystems, Germany). Coronal brain sections were permeabilized in 0.3% Triton X-100 (CAS No. 9002-93-1, United States Biological, USA) dissolved in 0.1 M phosphate buffer solution (PBTx, pH = 7.4) for 1 hr at room temperature. These sections were then subjected to a blocking step by incubating them in 10% horse serum (Catalog No. 16050122, Gibco, New Zealand) dissolved in 0.3% PBTx solution for 2 hrs at room temperature. To assess the expression of HA-tagged hM3Dq-DREADD with markers of excitatory neurons within the neocortex and the hippocampus, permeabilized and blocked brain tissue sections were incubated in a primary antibody cocktail containing rat anti-HA (1:200; Catalog No. 10145700, Roche Diagnostics, Switzerland) antibody with rabbit anti-CaMKII (pan) (1:200; Catalog No. 3362BC, Cell Signaling Technology, USA) antibody dissolved in 0.3% PBTx solution for 3 days at 4°C. Sections were then washed thrice with 0.3% PBTx solution for 15 minutes each, followed by incubation with a secondary antibody cocktail containing goat anti-rat IgG antibody conjugated to Alexa Fluor 488 (1:500; Catalog No. A-11008, Invitrogen, USA) with goat anti-rabbit IgG antibody conjugated to Alexa Fluor 568 (1:500; Catalog No. A-11011, Invitrogen, USA) dissolved in 0.3% PBTx solution for 2 hrs at room temperature. Following sequential washes with 0.3% PBTx solution, sections were mounted on a slide using VECTASHIELD Antifade Mounting Medium with DAPI (Catalog No. H-1200, Vector Laboratories, USA), and images were acquired using a laser scanning confocal microscope (FV1200; Olympus Life Science, Japan). To count the number of GFAP-positive cells within the hippocampus, permeabilized and blocked brain tissue sections were incubated with rabbit anti-glial fibrillary acidic protein (GFAP, 1:500; Catalog No. G9269, Sigma-Aldrich, USA) antibody dissolved in 0.3% PBTx solution for 16-18 hrs at room temperature. After three sequential washes of 15 minutes each in 0.1 M phosphate buffer (PB) solution, sections were incubated with the donkey anti-rabbit IgG antibody conjugated to Alexa Fluor 488 (1:500; Catalog No. A-21206, Invitrogen, USA) dissolved in 0.3% PBTx solution for 2 hrs at room temperature. Following sequential washes with 0.1 M PB solution, sections were mounted on a slide using VECTASHIELD Antifade Mounting Medium with DAPI (Catalog No. H-1200, Vector Laboratories, USA). To count the number of ALDH1L1-positive cells within the neocortex, antigen retrieval was performed by incubating coronal brain sections in 10 mM sodium citrate buffer solution (pH = 6.0) for 10 minutes at 90°C. Sections were then cooled down to room temperature and further washed with 0.3% PBTx solution twice for 5 minutes each at room temperature. These sections were subjected to a blocking step by incubating them in 5% horse serum dissolved in 0.3% PBTx solution for 1 hr at room temperature. The tissue sections were then incubated with rabbit anti-ALDH1L1 (1:500; Catalog No. ab87117, Abcam, UK) primary antibody and 2.5% horse serum dissolved in 0.3% PBTx solution for 16-18 hrs at room temperature. Washes, incubation with secondary antibody, and mounting were performed as mentioned for GFAP immunostaining.

#### Cell quantification

Slides were assigned a random code and quantification was conducted by an experimenter blind to the treatment groups. Images were acquired at 20X magnification using a widefield epifluorescence microscope (AxioObserver Z1; ZEISS, Germany) and a camera (AxioCam MRm; ZEISS, Germany). For the dorsal (from bregma AP: -1.70 mm to -2.30 mm) and ventral (from bregma AP: -2.92 mm to -3.40 mm) hippocampus, representative images from each of the hippocampal subfields—CA1, CA3, dentate gyrus (DG), and hilus were acquired for both hemispheres across 2-3 coronal sections (at least 300 *µ*m apart) per dorsal or ventral hippocampus per brain. For the neocortex, representative images from the primary somatosensory cortex barrel field (S1BF; from bregma AP: 0.38 mm to -1.58 mm) were acquired as mentioned for the hippocampus. For cell counting analysis, GFAP- and ALDH1L1-positive cells were manually counted using ImageJ (National Institutes of Health, USA). For the hippocampus, GFAP-positive cells within the layers, viz. *stratum lacunosum–moleculare, stratum radiatum, stratum pyramidale, and stratum oriens*, whereas for the somatosensory cortex, ALDH1L1-positive cells within layers II/III and V, were considered for quantification assays. Cells per mm^2^ (cells/mm^2^) were calculated for each measurement after dividing the cell numbers by the total area (mm^2^).

### Neurometabolic analysis with (^1^H-(^13^C)) NMR spectroscopy

To investigate the influence of chronic postnatal DREADD-mediated activation of CaMKIIα-positive neurons on adult glial metabolism within the forebrain, the concentration of ^13^C-labeled metabolites in neocortical and hippocampal tissues was estimated using (^1^H-(^13^C)) NMR spectroscopy following infusion of [2-^13^C]acetate. Vehicle- and CNO-treated bigenic CaMKIIα-tTA::tetO-hM3Dq adult male mice were deprived of food 6 hrs prior to cannulation of the tail vein to infuse [2-^13^C]acetate (1 mol/L, pH = 7.0; Acetic-2-^13^C acid sodium salt; Catalog No. CLM-381-10, Cambridge Isotope Laboratories, USA) dissolved with unlabeled glucose (0.225 mol/L) in deionized water. The [2-^13^C]acetate solution was administered as a bolus of 1.25 mmol/kg in 15 seconds, followed by an exponentially decreasing infusion rate for 2 minutes. Blood was withdrawn from the retro-orbital sinus before the end of the experiment while keeping the animal under mild isoflurane anesthesia. The collected blood was centrifuged to separate plasma, which was frozen in liquid nitrogen and stored at -80°C until further analysis. After exactly 10 minutes from the onset of [2-^13^C]acetate infusion, mice were euthanized by focused-beam microwave irradiation (FBMI; 4 kW for 0.94 seconds) using a microwave fixation system (MMW-05; Muromachi Kikai, Japan) to instantaneously arrest neurometabolic activity. The neocortical and hippocampal tissues were microdissected on ice, snap-frozen with liquid nitrogen, and stored at -80°C until further processing.

Metabolites were extracted from snap-frozen neocortical and hippocampal tissues as previously described (Patel *et al*., 2001). In brief, the snap-frozen tissues were homogenized in 0.1 N HCl dissolved in methanol to achieve a concentration of 3X volume/weight (v/w). [2-^13^C]Glycine (0.2 µmol; Catalog No. CLM-136, Cambridge Isotope Laboratories, USA) was added to the homogenate as an internal concentration reference. After homogenization, 6X (v/w) 90% ice-cold ethanol was added, and the tissue was further homogenized. The homogenate was then centrifuged at 16,000 *g* for 45 minutes at 4°C. The supernatant was passed through the Chelex column (Catalog No. 142-2822, Bio-Rad, USA), and the pH of the extract was adjusted to 7.0. The lyophilized extracts were dissolved in phosphate-buffered deuterium oxide (Catalog No. DLM-4-99-1000, Cambridge Isotope Laboratories, USA) containing sodium 3-trimethylsilyl[2,2,3,3-D4]-propionate (TSP, 0.25 mmol/L; Catalog No. 269913-1G; Sigma-Aldrich, USA), where TSP was used as a reference against which all chemical shifts were measured.

Blood plasma (100 *µ*L) was mixed with sodium formate (1 *µ*mol/L; Catalog No. 67253-1G; Sigma-Aldrich, USA) prepared in deuterium oxide (450 *µ*L) and passed through a 10 kDa cut-off centrifugal filter (Catalog No. 82031-350; VWR International, USA) by centrifugation at 13000 *g* for 90 minutes to remove macroparticles. (^1^H) NMR spectra were acquired using a 600 MHz spectrometer (AVANCE II; Bruker, Germany) keeping formate as an internal reference. The percent ^13^C labeling of acetate-C2 was calculated by dividing the area of ^13^C-coupled satellites by the total area (^12^C + ^13^C) observed at 1.9 ppm. The (^1^H-(^13^C)) NMR spectra of neocortical and hippocampal tissue extracts were acquired using a 600 MHz spectrometer as previously described (de Graaf *et al*., 2003; Bagga *et al*., 2013). In brief, two spin-echo (^1^H) NMR spectra were recorded in an OFF/ON ^13^C inversion pulse configuration. The free induction decays (FID) were zero-filled, apodized to Lorentzian line broadening, Fourier transformed, and phase-corrected. The ^13^C-edited NMR spectra were obtained by subtracting the sub-spectrum obtained with the ^13^C inversion pulse from the respective spectra acquired without the inversion pulse. The concentration of neurometabolites was estimated relative to [2-^13^C]glycine, which was added during extraction of metabolites from brain tissue. The ^13^C enrichment of various metabolites at different carbon positions was determined as the ratio of the peak areas in the (^1^H-(^13^C)) NMR difference spectrum (^13^C only) to the non-edited spectrum (^12^C + ^13^C), and was subsequently corrected for the natural abundance of ^13^C (1.1%).

#### Estimation of acetate oxidation metabolic rate

In the brain, acetate in the bloodstream is preferentially transported into astrocytes and oxidized therein (Patel *et al*., 2010). Oxidation of [2-^13^C]acetate in astrocytes labels astrocyte glutamine_C4_ (Gln_C4_) via the tricarboxylic acid (TCA) cycle, which is transported to glutamatergic and GABAergic neurons. In neurons, the ^13^C label from Gln_C4_ is transferred to glutamate_C4_ (Glu_C4_) and GABA_C2_ through neurotransmitter cycling between astrocytes and neurons (Patel *et al*., 2010) (**Figure S1**). Unlabeled glucose acts as a substrate for glycolysis in both neurons and astrocytes, which generates pyruvate that further acts as a substrate for the TCA cycle (**Figure S1**).

The metabolic rates of acetate oxidation following simultaneous [2-^13^C]acetate and unlabeled glucose infusion, indicative of astrocyte metabolism, were calculated using the estimated concentration of various amino acid metabolites labeled with ^13^C as previously described (Patel *et al*., 2005, 2018; Mishra *et al*., 2018).

The total cerebral metabolic rate of acetate oxidation (CMR_Ace(Ox)_ or MR_Glia_) was calculated using the following equation:

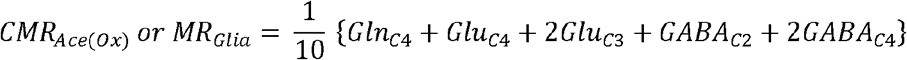

where Glu_Ci_, GABA_Ci_, Gln_Ci_, and Asp_Ci_ refer to the concentrations of ^13^C-labeled glutamate, GABA, glutamine, and aspartate, respectively, at the corresponding ‘i’th carbon following [2-^13^C]acetate and unlabeled glucose infusion.

### Statistical Analysis

Statistical analyses were performed using Prism 11 software (GraphPad Software Inc., USA). Unless specified, most experiments had two treatment groups (PNCNO treatment as the variable). To test the normality of distribution, the D’Agostino-Pearson normality test was performed. All experiments with data that fit the Gaussian distribution were subjected to a two-tailed, unpaired Student’s *t*-test. A Welch correction was applied when a significant difference in the variance between the groups was observed. Data that did not fit a Gaussian distribution were subjected to the Mann-Whitney *U* test.

To analyze longitudinal body weight differences across treatment (vehicle and PNCNO) and treatment window (P2 to P14), a two-way repeated measures ANOVA with Greenhouse-Geisser correction was performed. For gene expression data, an additional false discovery rate (FDR) analysis was carried out for calculated *p*-values using the Benjamini and Hochberg method, and these are denoted as FDR-corrected *p*-values. All data are represented as mean ± standard error of the mean (S.E.M.), and statistical significance was set at *p* < 0.05.

## RESULTS

### Selective expression of hM3Dq-DREADD in CaMKIIα-positive forebrain excitatory neurons in bigenic CaMKIIα-tTA::tetO-hM3Dq mice

To examine the consequences of chronic postnatal hM3Dq-DREADD-mediated activation of CaMKIIα-positive neurons on adult glial function and metabolism, bigenic CaMKIIα-tTA::tetO-hM3Dq mice were generated **(Figure 1A)**. The expression of the HA-tagged hM3Dq-DREADD was characterized in the forebrain, wherein the DREADD is expressed under the CaMKIIα-tTA promoter in forebrain excitatory neurons **(Figure 1B)**. Western blotting analysis confirmed the presence of HA protein within the neocortex and hippocampi in adult bigenic CaMKIIα-tTA::tetO-hM3Dq mice **(Figure 1C)**. Using immunofluorescence analysis, we also confirmed that expression of HA-tagged hM3Dq-DREADD colocalized with CaMKIIα-positive neurons, in both the neocortex **(Figure 1D)** and the hippocampus **(Figure 1E)**, of adult bigenic CaMKIIα-tTA::tetO-hM3Dq mice. A prior study from our lab extensively characterized HA-tagged hM3Dq-DREADD expression during the postnatal window (P7) and showed that chronic CNO administration in bigenic CaMKIIα-tTA::tetO-hM3Dq pups during the postnatal window evoked an increase in expression of immediate early genes, *viz*., *p-Erk/Erk* and *Cfos*, in the hippocampus compared to the vehicle treatment group (Pati *et al*., 2020). We have also previously established that bath application of CNO to acute hippocampal slices resulted in a hM3Dq-DREADD-mediated robust spiking activity of CA1 pyramidal neurons, as well as that hM3Dq-DREADD was not colocalized with PV-positive interneurons or GFAP-positive astrocytes within the forebrain, and brain regions like the hypothalamus and the pallidum, which lack the CaMKIIα promoter, did not express the HA-tagged hM3Dq-DREADD.

**Figure 1:**
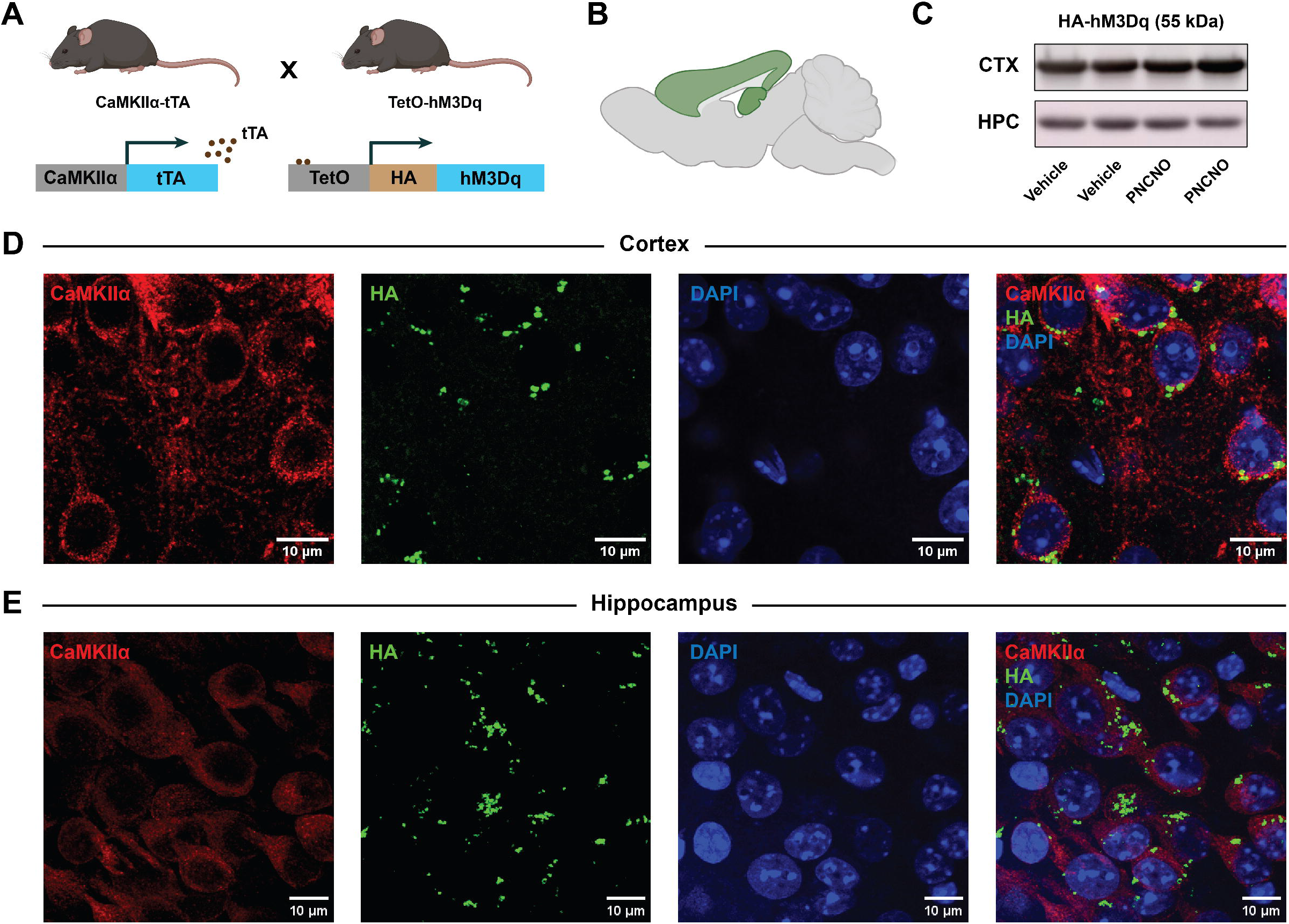
Selective expression of hM3Dq-DREADD in CaMKIIα-positive forebrain excitatory neurons in bigenic CaMKIIα-tTA::tetO-hM3Dq mice. **(A)** Shown is a schematic of the breeding strategy for the generation of the bigenic CaMKIIα-tTA::tetO-hM3Dq mouse line, wherein we selectively drive the expression of HA-tagged hM3Dq-DREADD in CaMKIIα-positive forebrain excitatory neurons. **(B)** Shown is a schematic of the forebrain—the region of HA-tagged hM3Dq-DREADD expression—through the sagittal view of a mouse brain. **(C)** Western blots detect the expression of HA-tag in the neocortex and the hippocampus, confirming the presence of HA-tagged hM3Dq-DREADD protein within the forebrain (n = 4 per brain region). **(D, E)** Shown are representative confocal images confirming the expression of HA-tag (green) within the neocortex **(D)** and the hippocampus **(E)**, along with colocalization of HA-tagged hM3Dq-DREADD with CaMKIIα-positive forebrain excitatory neurons (red) as identified with HA/CaMKIIα double immunofluorescence. Abbreviations: tTA – tetracycline transactivator; TetO – tetracycline operator; HA – hemagglutinin; CTX – cortex; HPC – hippocampus; PNCNO – postnatal clozapine N-oxide.

To assess the effect of CNO (1 mg/kg) administration in bigenic CaMKIIα-tTA::tetO-hM3Dq pups during the postnatal window on physiological growth, we recorded the average weight of pups in every litter across the vehicle and postnatal CNO (PNCNO) treatment groups **(Figure S2A)**. Oral administration of CNO, once a day from P2 to P14, did not alter the body weight of growing pups, measured across the treatment window **(Figure S2B)** or later in adulthood **(Figure S2C)**, compared to the vehicle-treated cohort. This is in agreement with our prior study, which noted that chronic CNO (1 mg/kg) administration from P2 to P14 does not alter normal physiological growth during the postnatal window in the PNCNO treatment group compared to the vehicle group (Pati *et al*., 2020).

Taken together, our observations from this work, along with our previous study, revealed that bigenic CaMKIIα-tTA::tetO-hM3Dq mice express the HA-tagged hM3Dq-DREADD in CaMKIIα-positive neurons within the neocortex and the hippocampus.

### Chronic postnatal chemogenetic activation of CaMKIIα-positive forebrain excitatory neurons increases anxiety-like behavior in adult male mice

We next assessed the behavioral consequences of chronic postnatal hM3Dq-mediated chemogenetic activation of CaMKIIα-positive forebrain excitatory neurons upon CNO administration. To do this, we subjected adult bigenic CaMKIIα-tTA::tetO-hM3Dq male mice with a history of either vehicle or PNCNO treatment to the elevated plus maze test (EPM) **(Figure 2A)**. We noted a significant increase in anxiety-like behavior on the EPM test in animals with a history of PNCNO treatment compared to the vehicle-treated controls **(Figure 2B)**, as shown with a significant decrease in percent time spent in the open arms of the EPM arena (*t*(18) = 2.215, *p* = 0.040) **(Figure 2C)**, as well as a trend towards a decrease in percent distance traveled in the open arms (*t*(18) = 1.856, *p* = 0.080) **(Figure 2D)**. The total distance traveled in the EPM arena was unchanged between the vehicle and PNCNO treatment groups **(Figure 2E)**, indicating that general locomotion was not impacted. Collectively, these observations indicate that chronic postnatal hM3Dq-mediated chemogenetic activation of forebrain excitatory neurons increases anxiety-like behavior in adult male mice. Our prior study showed that chronic hM3Dq-mediated activation of forebrain excitatory neurons during the postnatal window, but not juvenile or adult window, increases both anxiety- and despair-like behavior in bigenic CaMKIIα-tTA::tetO-hM3Dq adult male mice (Pati *et al*., 2020).

**Figure 2:**
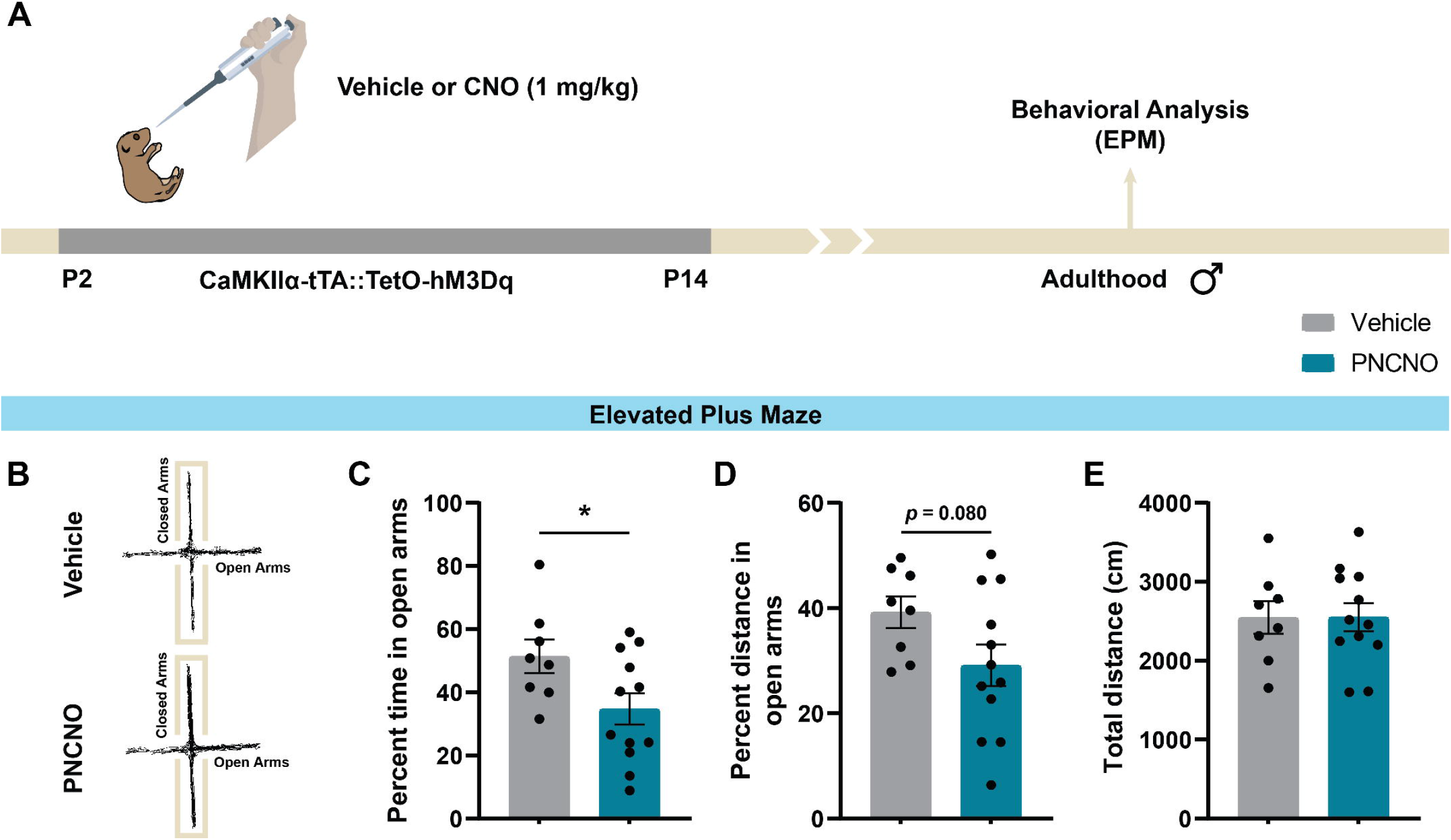
Chronic postnatal chemogenetic activation of CaMKIIα-positive forebrain excitatory neurons increases anxiety-like behavior in adult male mice. **(A)** Shown is a schematic of the experimental paradigm where bigenic CaMKIIα-tTA::tetO-hM3Dq mouse pups were orally administered vehicle or CNO (1 mg/kg) from P2 to P14 and then left undisturbed until adulthood prior to subjecting adult male mice to behavioral analysis on the EPM test. **(B)** Shown are representative tracks of vehicle-treated (upper panel) and PNCNO-treated (lower panel) bigenic CaMKIIα-tTA::tetO-hM3Dq adult male mice on the EPM test. Bigenic CaMKIIα-tTA::tetO-hM3Dq adult male mice with a history of PNCNO treatment exhibited an increased anxiety-like behavior on the EPM test, as evidenced by a significant decrease in percent time spent in open arms **(C)**, and a trend towards decreased percent distance traveled in the open arms **(D)**, compared to the vehicle-treated controls. **(E)** The total distance traveled was not significantly altered in PNCNO-treated adult male mice compared to vehicle-treated controls (n = 8 vehicle and 12 PNCNO). Results are expressed as the mean ± S.E.M. \**p* < 0.05, compared to vehicle-treated controls using the two-tailed, unpaired Student’s *t*-test. Abbreviations: PNCNO – postnatal clozapine N-oxide; EPM – elevated plus maze.

### Chronic postnatal chemogenetic activation of CaMKIIα-positive forebrain excitatory neurons alters glial marker and glia-associated expression in the neocortex and hippocampi of adult male mice

Given prior evidence indicating glial dysfunction associated with early adversity-evoked emergence of anxio-depressive behaviors (Banasr *et al*., 2008; Rajkowska & Miguel-Hidalgo, 2008; Abbink *et al*., 2019; Teissier *et al*., 2019), and our current observations indicating that chronic chemogenetic activation of CaMKIIα-positive forebrain excitatory neurons enhanced anxiety-like behavior in adulthood, we investigated the consequence of chronic postnatal CNO-mediated DREADD activation of CaMKIIα-positive forebrain excitatory neurons on glial function in adulthood. To examine this, we employed complementary approaches to measure relative gene and protein expression of glial and glia-associated markers within the neocortex and hippocampi of bigenic CaMKIIα-tTA::tetO-hM3Dq adult male mice with a history of either vehicle or PNCNO treatment **(Figure 3A)**. We performed qPCR analysis to assess gene expression of the glial fibrillary acidic protein (*Gfap*), a major marker of reactive astrocytes; aldehyde dehydrogenase 1, member L1 (*Aldh1l1*), a marker of both non-reactive and reactive astrocytes; S100 calcium binding protein beta (*S100β*); vimentin, an intermediate filament protein that labels immature glia and is enhanced in reactive astrocytes; excitatory amino acid transporters (*Eaat*); and GABA transporters (*Gat*) that mediate neurotransmitter cycling between astrocytes and neurons. In the cortex, we observed a significant decline in the relative gene expression of *Gfap* (*t*(15) = 3.290, *p* = 0.005), *Aldh1l1* (*t*(15) = 2.139, *p* = 0.049), *S100β* (Mann-Whitney *p* = 0.005), *Vimentin* (*t*(15) = 2.153, *p* = 0.048), *Eaat1* (*t*(15) = 2.671, *p* = 0.017), and *Eaat3* (Mann-Whitney *p* = 0.008), along with a trend towards significance for *Gat2* (*t*(15) = 1.872, *p* = 0.081) in adult male mice with a history of PNCNO treatment **(Figure 3B)**. In contrast, in the hippocampus, we observed a significant upregulation of relative gene expression of *Gfap* (*t*(16) = 4.201, *p* = 0.0007), *Aldh1l1* (*t*(16) = 3.950, *p* = 0.001), *Eaat1* (Mann-Whitney *p* = 0.012), *Eaat2* (*t*(11.82) = 4.587, *p* = 0.0006, Welch’s correction), *Eaat3* (*t*(16) = 5.257, *p* < 0.0001), *Gat1* (*t*(13) = 4.727, *p* = 0.0004, Welch’s correction), and *Gat3* (*t*(15) = 4.317, *p* = 0.0006), as well as a trend towards significance for *Eaat4* (*t*(16) = 1.891, *p* = 0.077) in PNCNO-treated adult male mice **(Figure 3C)**.

**Figure 3:**
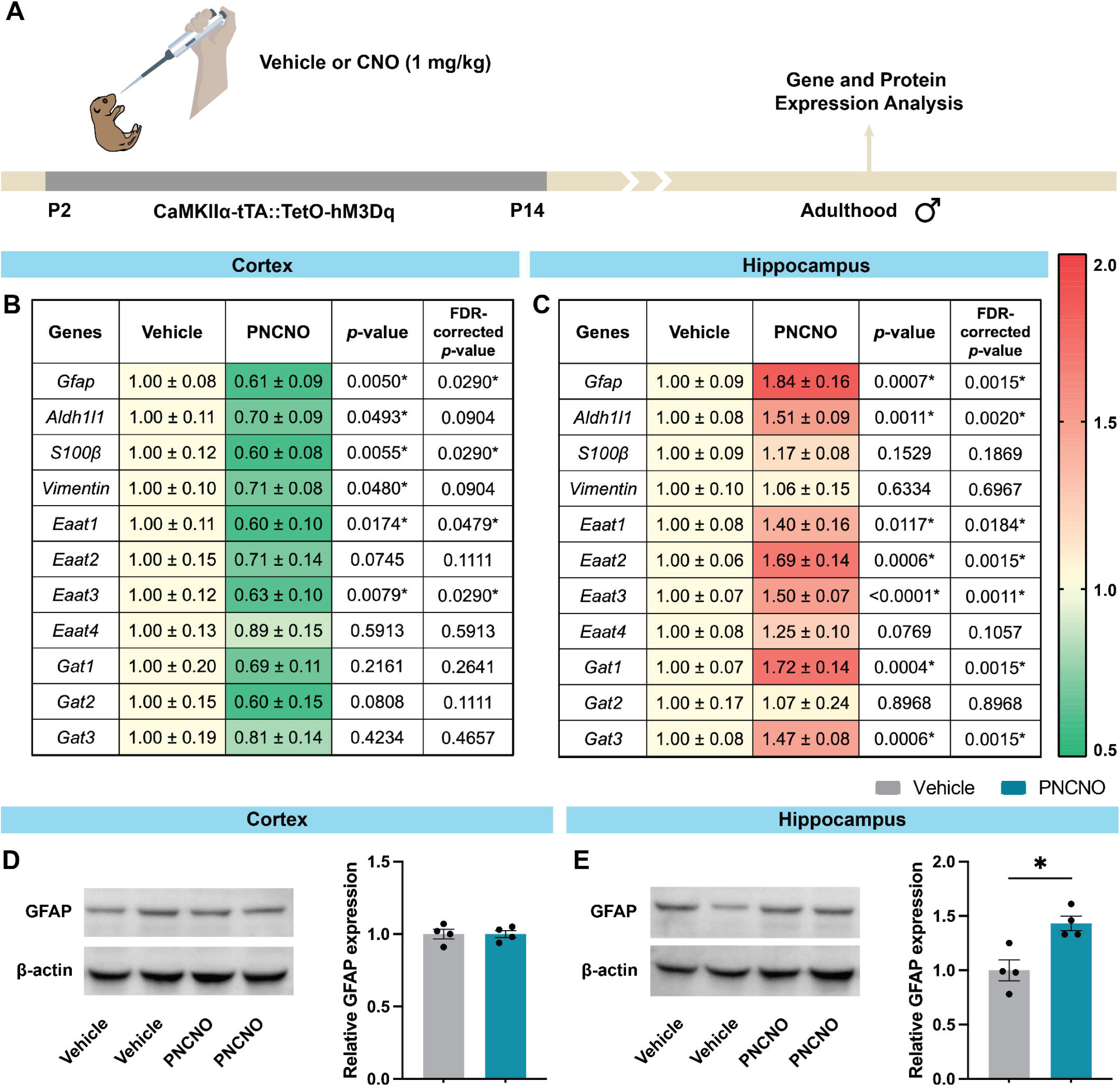
Chronic postnatal chemogenetic activation of CaMKIIα-positive forebrain excitatory neurons alters glial marker and glia-associated expression in the neocortex and hippocampi of adult male mice. **(A)** Shown is a schematic of the experimental paradigm where bigenic CaMKIIα-tTA::tetO-hM3Dq mouse pups were orally administered vehicle or CNO (1 mg/kg) from P2 to P14 and then left undisturbed until adulthood prior to subjecting adult male mice to gene and protein expression analysis using qPCR and western blotting, respectively. **(B, C)** Shown are normalized gene expression levels for glial and glia-associated markers in the neocortex **(B)** and the hippocampus **(C)** of PNCNO-treated adult male mice, represented as fold-change of vehicle-treated controls (1.00 ± S.E.M.) (n = 8-10 per group). Heat maps indicate the extent of gene regulation. Also shown are *p*-values and FDR-corrected *p*-values for the statistical tests performed for each gene. **(D, E)** Shown are Western blots of GFAP and β-actin (left) and graphs of relative protein expression of GFAP (normalized to β-actin), represented as fold-change of vehicle-treated controls (right, 1.00 ± S.E.M.) in the neocortex **(D)** and the hippocampus **(E)** (n = 4 per group for both cortex and hippocampus). Results are expressed as the mean ± S.E.M. \**p* < 0.05, compared to vehicle-treated controls using the two-tailed, unpaired Student’s *t*-test, Welch’s *t*-test, or the Mann-Whitney *U* test. Abbreviations: PNCNO – postnatal CNO; qPCR – quantitative polymerase chain reaction; FDR – false discovery rate; GFAP – glial fibrillary acidic protein; ALDH1L1 – aldehyde dehydrogenase 1 family member L1; S100β – S100 calcium-binding protein β; EAAT – excitatory amino acid transporter; GABA – γ-aminobutyric acid; GAT – GABA transporter type 1.

We next sought to measure the relative protein expression of GFAP within the neocortex and hippocampi derived from vehicle- and PNCNO-treated bigenic CaMKIIα-tTA::tetO-hM3Dq adult male mice with Western blotting analysis. We noted a region-specific alteration in GFAP protein expression levels, with a significant increase in the hippocampus (*t*(6) = 3.692, *p* = 0.01) **(Figure 3E)**, but no statistically significant change in GFAP protein expression in the neocortex **(Figure 3D)** of PNCNO-treated mice, compared to the vehicle-treated cohort. Collectively, our findings reveal that chronic chemogenetic activation of forebrain excitatory neurons in postnatal life can program significant bidirectional alterations in gene expression of multiple markers associated with astrocyte structure and function, including changes in transporters that modulate neurotransmitter fluxes between astrocytes and neurons.

### Chronic postnatal chemogenetic activation of CaMKIIα-positive forebrain excitatory neurons does not alter astrocyte number in the cortex and hippocampi of adult male mice

We examined whether our gene and protein expression data correlated with any change in astrocyte counts within the neocortex and the hippocampus. To assess this, we performed a cell quantification analysis of ALDH1L1 immunoreactive cells within the primary somatosensory cortex (as a representative of the neocortex), and GFAP immunoreactive cells within the dorsal and ventral hippocampus **(Figure 4A)**. We chose to analyze astrocyte numbers separately in the dorsal and ventral hippocampus because of the inherent functional distinctions between these two regions, with the dorsal hippocampus implicated in cognitive functions and the ventral hippocampus being implicated in the regulation of stress, emotion, and affect (Fanselow & Dong, 2010; Lee *et al*., 2017, 2019). Almost all astrocytes in the hippocampus are GFAP-positive; hence, we counted GFAP-positive cells to estimate the astrocyte number in the hippocampus. However, in the neocortex, GFAP is only expressed by white matter astrocytes, and not grey matter astrocytes (Walz, 2000). Hence, we counted ALDH1L1-positive cells to estimate astrocyte number in the neocortex, as ALDH1L1 is a specific pan-astrocyte marker that is expressed in all cortical astrocytes (Yang *et al*., 2011).

**Figure 4:**
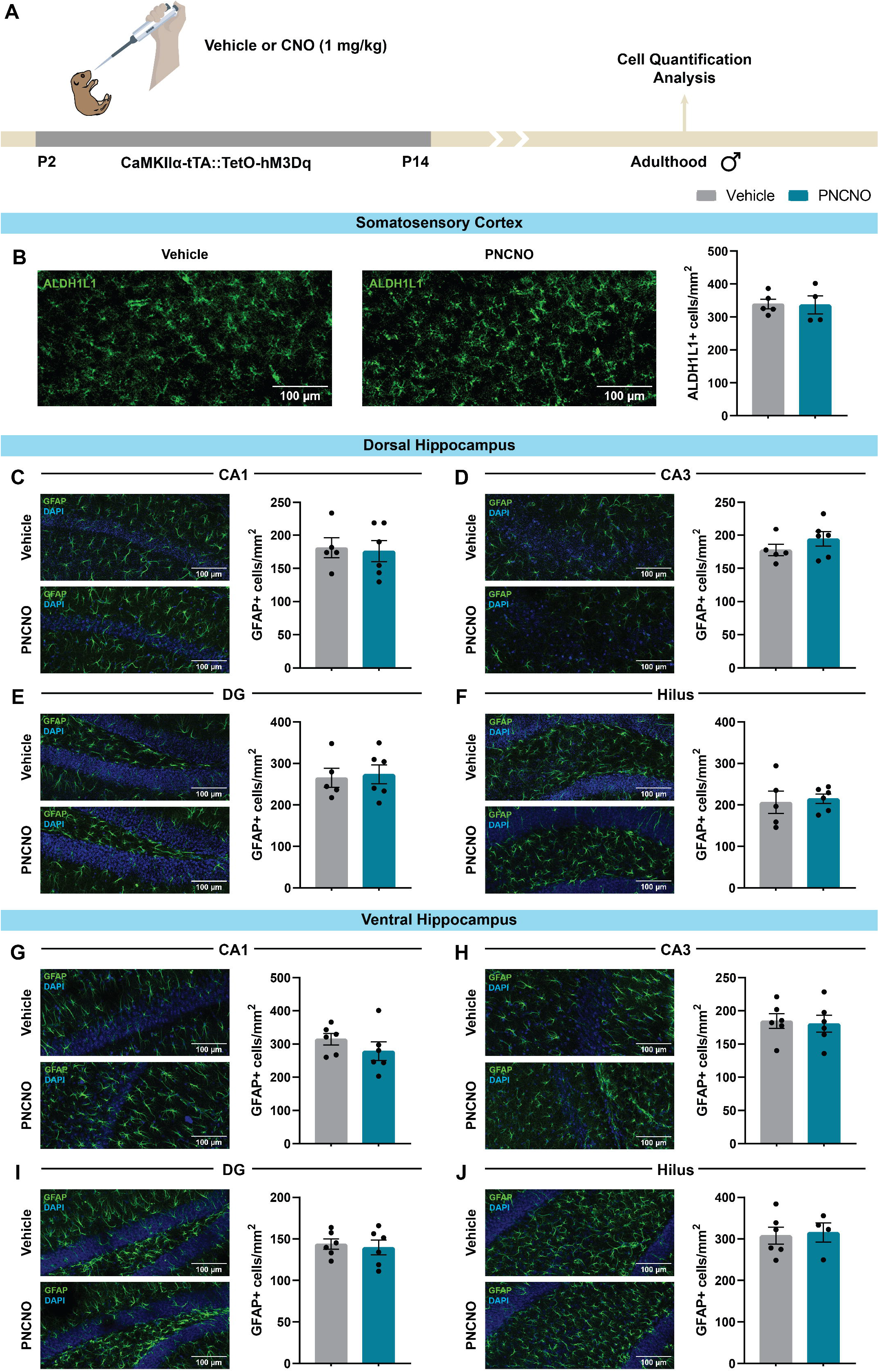
Chronic postnatal chemogenetic activation of CaMKIIα-positive forebrain excitatory neurons does not alter astrocyte number in the cortex and hippocampi of adult male mice. **(A)** Shown is a schematic of the experimental paradigm where bigenic CaMKIIα-tTA::tetO-hM3Dq mouse pups were orally administered vehicle or CNO (1 mg/kg) from P2 to P14 and then left undisturbed until adulthood prior to subjecting adult male mice to cell counting analyses of ALDH1L1- and GFAP-positive astrocytes within the neocortex and hippocampi, respectively. Cell counting analysis of ALDH1L1-positive cells within the S1BF cortex **(B)** revealed that chronic postnatal chemogenetic activation of forebrain excitatory neurons does not alter astrocyte cell numbers (ALDH1L1-positive cells/mm^2^) in PNCNO-treated adult male mice compared to vehicle-treated control male mice (n = 5 vehicle and 4 PNCNO). Similarly, no significant differences in astrocyte cell numbers (GFAP-positive cells/mm^2^) were observed within the dorsal hippocampal **(C, D, E, F)** and ventral hippocampal **(G, H, I, J)** subfields, *viz*., dorsal and ventral CA1 **(C, G)**, CA3 **(D, H)**, DG **(E, I)**, as well as hilus **(F, J)**, in PNCNO-treated adult male mice compared to vehicle-treated control male mice (n = 5 vehicle and 6 PNCNO for dorsal hippocampus; n = 4-6 per group for ventral hippocampus). Results are expressed as the mean ± S.E.M., and groups are compared using the two-tailed, unpaired Student’s *t*-test. Abbreviations: PNCNO – postnatal CNO; ALDH1L1 – aldehyde dehydrogenase 1 family member L1; GFAP – glial fibrillary acidic protein; S1BF – primary somatosensory barrel field cortex; CA – *Cornu Ammonis*; DG – dentate gyrus.

In the primary somatosensory barrel field cortex (S1BF), the number of ALDH1L1-positive astrocytes was unaltered in bigenic CaMKIIα-tTA::tetO-hM3Dq adult male mice with a history of PNCNO treatment compared to vehicle-treated controls **(Figure 4B)**. In the dorsal hippocampus, the astrocyte number within hippocampal subfields, namely, CA1 **(Figure 4C)**, CA3 **(Figure 4D)**, dentate gyrus (DG) **(Figure 4E)**, and hilus **(Figure 4F)** was unchanged between vehicle- and PNCNO-treated adult male mice, as evident by no alterations in the number of GFAP-positive cells per mm^2^ within these hippocampal subfields. In the ventral hippocampus, we noted a similar observation where the astrocyte numbers within hippocampal subfields, namely, CA1 **(Figure 4G)**, CA3 **(Figure 4H)**, dentate gyrus (DG) **(Figure 4I)**, and hilus **(Figure 4J)**, were unchanged between vehicle- and PNCNO-treated adult male mice. Taken together, our results indicate that chronic postnatal chemogenetic activation of forebrain excitatory neurons does not alter astrocyte numbers in the somatosensory cortex as well as the dorsal and ventral hippocampus of PNCNO-treated bigenic CaMKIIα-tTA::tetO-hM3Dq adult male mice.

### Chronic postnatal chemogenetic activation of CaMKIIα-positive forebrain excitatory neurons results in long-lasting alterations in neurotransmitter flux and glial metabolic rate in the neocortex and hippocampi of adult male mice

Preclinical and clinical studies indicate dysregulation of forebrain glutamatergic and GABAergic neurotransmission as an important contributor to the pathophysiology of mood-related disorders, such as anxiety, depression, and schizophrenia (Choudary *et al*., 2005; Kendell *et al*., 2005; Sanacora *et al*., 2012; Duman *et al*., 2019). Metabolic dysfunction of glutamatergic and GABAergic systems has been suggested to be an important endophenotype for these mood-related disorders (Hasler & Northoff, 2011; Veeraiah *et al*., 2014; Godfrey *et al*., 2018; Sekar *et al*., 2019). Given our qPCR analyses reveal a profound dysregulation of glutamate and GABA transporters as a consequence of chronic postnatal DREADD-mediated activation of forebrain excitatory neurons, we hypothesized that such altered expression may reveal metabolic dysfunction of glutamatergic and GABAergic systems. To study this, we performed metabolic analysis within the neocortex and hippocampus derived from bigenic CaMKIIα-tTA::tetO-hM3Dq adult male mice with a history of either vehicle or PNCNO treatment using an NMR tracing approach involving the infusion of [2-^13^C]acetate and unlabeled glucose **(Figure 5A)**. This approach is based on the phenomenon of preferential transport of acetate present in the bloodstream into astrocytes (Patel *et al*., 2010). [2-^13^C]Acetate is preferentially oxidized within astrocytes and metabolized to astrocytic glutamine_C4_ (Gln_C4_) via the TCA cycle. Unlabeled glucose is metabolized to pyruvate via glycolysis in both neurons and astrocytes. Pyruvate is further oxidized to other metabolites via the TCA cycle. In a three-compartment metabolic model **(Figure S1)**, Gln_C4_ is transported to glutamatergic and GABAergic neurons, where it transfers the ^13^C label to glutamate_C4_ (Glu_C4_) or GABA_C2_ via the glutamine-glutamate or glutamine-GABA cycle, respectively, between astrocytes and neurons (Patel *et al*., 2010; Saba *et al*., 2017). The metabolic rate of acetate oxidation in astrocytes was estimated within the neocortex and the hippocampus using the three-compartment metabolic rate model. We did not observe any significant differences in the levels of glutamate, GABA, glutamine, aspartate, N-acetylaspartate, alanine, lactate, inositol, taurine, choline and creatine from non-edited (^1^H-(^12^C + ^13^C)) NMR spectra measured using [2-^13^C]glycine as reference within the neocortex and hippocampus of PNCNO-treated bigenic CaMKIIα-tTA::tetO-hM3Dq adult male mice compared to their vehicle-treated cohort **(Figure S3)**.

**Figure 5:**
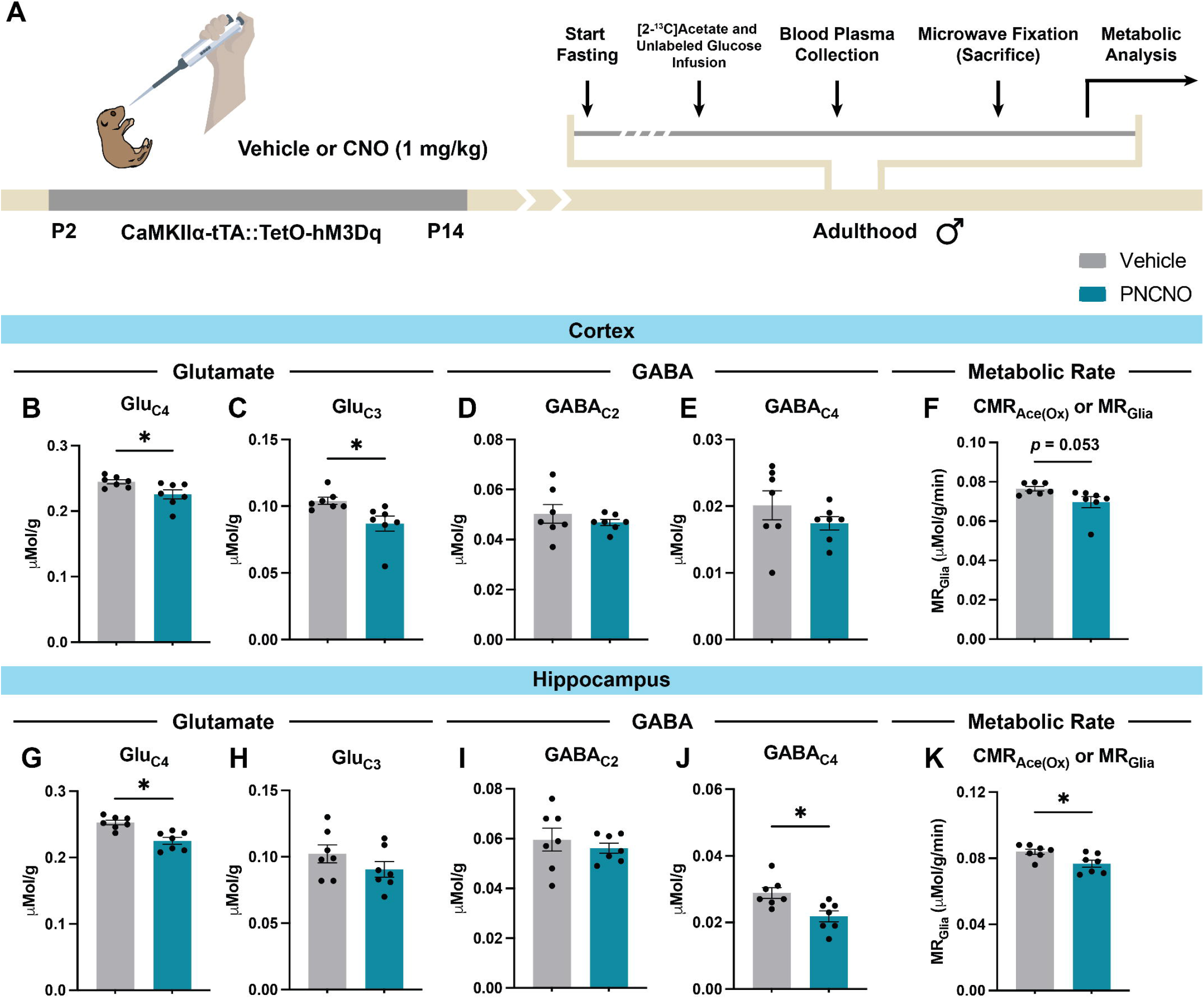
Chronic postnatal chemogenetic activation of CaMKIIα-positive forebrain excitatory neurons results in long-lasting alterations in neurotransmitter flux and glial metabolic rate in the neocortex and hippocampi of adult male mice. **(A)** Shown is a schematic of the experimental paradigm where bigenic CaMKIIα-tTA::tetO-hM3Dq mouse pups were orally administered vehicle or CNO (1 mg/kg) from P2 to P14 and then left undisturbed until adulthood prior to subjecting adult male mice to metabolic analysis using (^1^H-(^13^C)) NMR spectroscopy. (^1^H-(^13^C)) NMR spectra of neocortical and hippocampal tissue extracts were recorded and analyzed as previously described (Patel et al., 2001; Saba et al., 2017). Briefly, adult male mice were subjected to fasting for 6 hrs, following which [2-^13^C]acetate and unlabeled glucose were infused via the tail vein. Blood plasma was collected via retro-orbital bleeding, and mice were sacrificed after 10 min following acetate and glucose infusion by focused-beam microwave irradiation to instantly arrest neurometabolic activity. Bigenic CaMKIIα-tTA::tetO-hM3Dq adult male mice with a history of PNCNO treatment showed a significant decrease in ^13^C-labeled neurotransmitter turnover, as indicated by a decline in the concentration of Glu_C4_ **(B)** and Glu_C3_ **(C)** within the neocortex, as well as Glu_C4_ **(G)** and GABA_C4_ **(J)** within the hippocampus, compared to the vehicle-treated control male mice. The concentration of ^13^C-labeled GABA_C2_ **(D)** and GABA_C4_ **(E)** within the neocortex, as well as Glu_C3_ **(H)** and GABA_C2_ **(I)** within the hippocampus, was unaltered across treatment groups. The cerebral metabolic rate of astrocyte-mediated acetate oxidation (CMR_Ace(Ox)_), indicative of astrocyte metabolism (MR_Glia_), was also decreased in the neocortex **(F)** and the hippocampus **(K)** of PNCNO-treated adult male mice compared to the vehicle-treated controls (n = 7 per group). Results are expressed as the mean ± S.E.M. \**p* < 0.05, compared to vehicle-treated controls using the two-tailed, unpaired Student’s *t*-test or Welch’s *t*-test. Abbreviations: PNCNO – postnatal CNO; Glu – glutamate; GABA – γ-aminobutyric acid; CMR – cerebral metabolic rate; NMR – nuclear magnetic resonance.

We then set out to estimate the concentration of glutamate and GABA metabolites carrying the ^13^C-label from [2-^13^C]acetate at various carbon positions from the ^13^C-edited (^1^H-(^13^C)) NMR spectra of neocortical and hippocampal tissue extracts derived from the vehicle and PNCNO treatment groups. In the neocortex, we noted a significant reduction in glutamate labeling from [2-^13^C]acetate as revealed by a significant decrease in the concentration of ^13^C-labeled Glu_C4_ (*t*(12) = 2.527, *p* = 0.027) **(Figure 5B)** and Glu_C3_ (*t*(12) = 2.734, *p* = 0.018) **(Figure 5C)** in the PNCNO treatment group. We did not note any difference in GABA_C2_ **(Figure 5D)** and GABA_C4_ **(Figure 5E)** in the neocortex. In the hippocampus, both glutamate and GABA labeling were altered, as evidenced by a significant reduction in the concentration of ^13^C-labeled Glu_C4_ (*t*(12) = 4.351, *p* = 0.001) **(Figure 5G)** and GABA_C4_ (*t*(12) = 3.047, *p* = 0.010) **(Figure 5J)** in the PNCNO treatment group. However, we did not observe any change in the concentration of ^13^C-labeled Glu_C3_ **(Figure 5H)** and GABA_C2_ **(Figure 5I)** in the hippocampus.

Further, we sought to determine the cerebral metabolic rate of acetate oxidation in astrocytes within the neocortex and the hippocampus from the concentration of ^13^C-labeled amino acids as previously described (Patel *et al*., 2005; Mishra *et al*., 2018). We observed an overall reduction in the rate of acetate oxidation, indicative of astrocyte metabolism, within the neocortex (*t*(7) = 2.276, *p* = 0.053, Welch’s correction) **(Figure 5F)** and the hippocampus (*t*(12) = 2.702, *p* = 0.019) **(Figure 5K)** of PNCNO-treated bigenic CaMKIIα-tTA::tetO-hM3Dq adult male mice compared to their vehicle-treated controls.

Collectively, our findings suggest that chronic postnatal chemogenetic activation of CaMKIIα-positive forebrain excitatory neurons results in a long-lasting decrease of astrocyte metabolic activity, along with glutamate and GABA labeling—indicating a persistent alteration in glutamatergic and GABAergic neurotransmission—within the forebrain in adulthood.

## DISCUSSION

Our study examines the influence of postnatal hM3Dq-mediated chemogenetic activation of CaMKIIα-positive forebrain excitatory neurons in programming adult anxiety-like behavior, and influencing astrocyte gene expression and function. We show that this transient chemogenetic perturbation during the postnatal window is sufficient to program increased anxiety-like behaviors, in keeping with our prior study (Pati *et al*., 2020). This was accompanied by brain region-specific transcriptional dysregulation of glial markers and glia-associated transporters in adulthood, with a decline in the neocortex and enhanced expression in the hippocampus, and an increase in relative GFAP protein expression in the hippocampus, but not the neocortex. Astrocyte cell numbers did not change within these forebrain regions. Interestingly, this chemogenetic perturbation was also associated with impaired astrocyte metabolism in the neocortex and hippocampus in adulthood: glutamate turnover declined in both the neocortex and hippocampus; GABA turnover declined in the hippocampus. This was accompanied by a significant reduction in overall metabolic rate of acetate oxidation within the hippocampus, with a trend towards a similar decline noted in the neocortex. Together, these findings reveal that chronic postnatal chemogenetic activation of forebrain excitatory programs long-lasting perturbations of both excitatory and inhibitory neurotransmission, as well as disrupting astrocyte-specific metabolism, suggesting compromised astrocyte-neuron metabolic coupling, which could perturb neurotransmitter homeostasis and synaptic function.

Our key focus in this study was on uncovering the impact of transient postnatal chemogenetic perturbation of forebrain excitatory neurons on astrocyte function and metabolism in adulthood. A substantial body of preclinical and clinical literature indicates that exposure to early-life stress alters astrocyte function in a stressor- and brain region-specific manner, with effects emerging at distinct temporal epochs following stressor exposure. For instance, hippocampal GFAP reactivity declines within 24 hrs after maternal deprivation (Saavedra *et al*., 2018; Réus *et al*., 2019), but is upregulated within a few days to weeks (Llorente *et al*., 2009; Réus *et al*., 2019). Whereas, maternal separation increases in astrocyte-specific EAAT1 and EAAT2 within the hippocampus (Martisova *et al*., 2012); however, limited nesting and bedding decreases EAAT1 in the paraventricular nucleus of the hypothalamus (Gunn *et al*., 2013), highlighting the distinct consequences of different early life stress models on astrocyte-associated measures. In our study, transient postnatal chemogenetic stimulation of forebrain excitatory neurons phenocopies specific anxiogenic behavioral consequences of early-life stress models, and we note that *Gfap* gene expression is bidirectionally altered, with a decrease in the neocortex and an increase in the hippocampus, and this trend was mimicked in terms of GFAP protein expression in the hippocampus, but not in the cortex. This regional specificity likely reflects differential sensitivity of the cortex and the hippocampus to perturbations in neuronal activity during early development. Given that we noted collective shifts in gene expression across multiple astrocyte-associated markers (*Gfap, Aldh1l1, Eaat*, and *Gat*), we sought to ask if astrocyte numbers are altered within the neocortex and hippocampus. Intriguingly, despite significant molecular changes, astrocyte cell numbers were unchanged in the primary somatosensory cortex, and dorsal and ventral hippocampi.

A key function of astrocytes is the regulation of glutamate and GABA cycling, which goes awry in mood-related disorders associated with early adversity (Sanacora & Banasr, 2013; Abbink *et al*., 2019; Çalışkan *et al*., 2020; Cathomas *et al*., 2022). Astrocytic metabolic dysfunction itself is also a core feature of clinical neuropsychiatric illnesses, and magnetic resonance spectroscopy (MRS) studies, especially ^1^H-MRS, suggest that the levels of key metabolites shuttled between neurons and astrocytes are altered in patients suffering from generalized anxiety and major depression, compared to healthy controls (Sanacora *et al*., 2004; Valentine & Sanacora, 2009; Dindarian *et al*., 2026). This is associated with dysregulated gene expression of EAAT1 and EAAT2, as well as glutamatergic and GABAergic transmitter systems in major depression (Choudary *et al*., 2005). Although there are plenty of ^1^H-MRS studies indicating perturbed overall metabolism associated with several neuropsychiatric disorders, we lack an understanding of changes in astrocyte-specific metabolic dynamics in the forebrain. Therefore, we capitalized on (^1^H-(^13^C)) NMR spectroscopy to study astrocyte-mediated neurotransmitter turnover and metabolism following [2-^13^C]acetate infusion. We find that enhancing DREADD-mediated Gq-signaling during early life has persistent effects on glutamate and GABA turnovers, and also impairs astrocyte-mediated metabolism in adult neocortex and hippocampus, broadly consistent with prior preclinical models and clinical studies of anxiety and depression (Veeraiah *et al*., 2014; Duman *et al*., 2019). It is also important to note that, while we see a brain region-specific decrease in astrocyte-mediated metabolism, the total concentration of key neurometabolites, such as N-acetylaspartate (NAA), choline, and creatine, were unaltered. This differs from prior studies, which noted broad brain region-dependent shifts in levels of these metabolites across multiple neuropsychiatric disorders, including major depression and generalized anxiety disorder (Dindarian *et al*., 2026). Given that our measurements were bulk across the entire neocortex, any potential regional changes may have been masked. Finally, a reduction in adult hippocampal astrocyte metabolic rate contrasts with enhanced neuronal metabolic rate noted previously following [1,6-^13^C_2_]glucose administration in mice using a similar chemogenetic treatment (Pati *et al*., 2020). These observations suggest a long-lasting bidirectional change in astrocytic versus neuronal metabolic activity in adulthood as a consequence of chronic enhanced Gq-coupled signaling in forebrain excitatory neurons during the postnatal window, pointing to the possibility of altered neurotransmitter recycling between forebrain neurons and astrocytes, that last long after the transient perturbation of Gq-signaling in forebrain excitatory neurons during postnatal life.

Although driving Gq signaling in the forebrain excitatory neurons impairs adult astrocyte function and metabolism, it is unclear whether these astrocyte-specific alterations directly contribute to the pathogenesis of early adversity-evoked psychopathology, or whether these maladaptive changes play a key role in contributing to the persistent effects on mood-related behavior. Models of early adversity program persistent structural and functional changes, particularly within the hippocampus and prefrontal cortex, which contribute to altered top-down control of anxiety-like behavior and stress response axes. Our work motivates future studies to address whether early life perturbations mediate some of the long-term mood-related behavioral changes via perturbation of astrocyte function and metabolism. Thus far studies have largely addressed consequences of perturbing Gq signaling in CaMKIIα-positive forebrain excitatory neurons and PV-interneurons, both of which evoke long-lasting changes in anxiety-related behaviors (Pati *et al*., 2020; Banerjee *et al*., 2022). It would be of interest to address whether chemogenetic perturbation of Gq signaling in astrocytes itself during postnatal life can also produce persistent changes in mood-related behaviors in adulthood. Despite having a limited understanding of the contribution of glial populations in the programming of anxio-depressive behavior, it is known that both astrocytes and microglia exhibit diverse changes based on the onset, severity, type, and duration of stressors during early life (Yamawaki *et al*., 2018; Banqueri *et al*., 2019; Réus *et al*., 2019; Abbink *et al*., 2020). While our study highlights the possible role of enhanced Gq signaling in early life on the emergence of mood-related disorders, as well as associated glial dysfunction and perturbed astrocyte metabolism, we lack the resolution of when and how these phenotypes occur. This gap can be bridged by a systematic study looking into the immediate, early, and intermediate effects of early adversity on glial function and metabolism. Having this temporal resolution would help determine whether glial changes precede, coincide with, or follow the emergence of behavioral abnormalities, and potentially how they map onto changes in the neuronal populations. Previous studies hinted that glial populations, especially microglia and astrocytes, may be amongst the first responders to early stressors, with neuronal phenotypes emerging later (Miguel-Hidalgo *et al*., 2000; Abbink *et al*., 2019; Guayasamin *et al*., 2025). Our findings raise an interesting possibility that perturbation of Gq signaling in forebrain excitatory neurons during critical periods can induce persistent astrocyte dysfunction, revealing that altered neuron-astrocyte cross-talk is associated with the programming of long-lasting effects on mood-related behaviors.

An important caveat is that our study only involves bigenic CaMKIIα-tTA::tetO-hM3Dq adult male mice with no experiments performed in adult female mice, in part due to the large numbers of bigenic mice required to be maintained for these experiments. Our prior study showed that chronic postnatal hM3Dq-DREADD-mediated activation of CaMKIIα-positive forebrain excitatory neurons increases anxiety-like, but not despair-like behavior in bigenic CaMKIIα-tTA::tetO-hM3Dq adult female mice (Pati *et al*., 2020), indicating that the effects of PNCNO treatment in these bigenic animals exhibit some sex differences. Further experimentation is needed to carefully analyze if hM3Dq chemogenetic activation of forebrain excitatory neurons during the postnatal window of life has any sex-dependent effects on adult glial function and metabolism. It is also important to note that, while we characterized astrocytic function using multiple complementary approaches, we did not assess changes in astrocyte morphology or ultrastructure features, which may have provided additional insights into functional consequences that we noted. Further studies are needed to address these open questions. In this regard, our chemogenetic strategy can augment naturalistic models of early-life adversity, such as maternal separation and maternal immune activation, to specifically understand the contribution of perturbed Gq-coupled GPCR signaling in early-life adversity-evoked emergence of anxio-depressive behaviors.

Taken together, our study demonstrates that chemogenetically enhancing Gq signaling in forebrain excitatory neurons during the postnatal window induces persistent, brain region-specific alterations in glial function and astrocyte metabolism. These glial phenotypes emerge in the absence of changes in astrocyte cell numbers in the neocortex, specifically within the primary somatosensory cortex and the hippocampus. Our findings underscore the persistent shift that emerges in astrocyte function and metabolism, suggestive of altered neuron-astrocyte cross-talk, following transient perturbations in early postnatal life that program long-lasting changes in adult anxiety-like behavior.

## ACKNOWLEDGEMENTS & FUNDING STATEMENT

This study is supported by intramural funds from the Department of Atomic Energy, Government of India, to the Tata Institute of Fundamental Research (TIFR) (RTI4003), and by JC Bose Fellowship (JCB/2021/000014) from the Anusandhan National Research Foundation (ANRF) and the Sree Ramakrishna Paramhamsa Research Grant (SreePVF/G/BS/19/1) from the Sree Padmavathi Venkateswara Foundation (SreePVF), Vijayawada, India, to Prof. Vidita Vaidya.

We thank Dr. Shital Suryavanshi, Dr. Archana Iyer, and the animal house staff at Tata Institute of Fundamental Research (TIFR), Mumbai, for their technical assistance. Schematics and illustrations used in this manuscript were created using BioRender.com.

## ETHICS STATEMENT

All experimental procedures were performed per the guidelines of the Committee for Control and Supervision of Experiments on Animals (CCSEA), Government of India, and were approved by the Tata Institute of Fundamental Research (TIFR/IAEC/2019-4) and the Centre for Cellular and Molecular Biology (CCMB/IAEC-32/2018) Institutional Animal Ethics Committees (IAEC). Care was taken across all experiments to minimize animal suffering and restrict the number of animals used.

## DATA AVAILABILITY STATEMENT

All data reported in this study will be available from the corresponding author upon request.

## AUTHOR CONTRIBUTIONS

Conceptualization: VV, SP

Data curation: AP, SP

Formal analysis: AP

Funding acquisition: VV, ABP

Investigation: AP, SP, KS, PT, PRC, DK, AB

Methodology: AP, SP, KS, PT, PRC, DK, AB, ABP, VV

Project Administration: VV

Resources: VV, ABP

Software: not applicable

Supervision: VV, ABP

Validation: SP, AP, KS, PT, PRC, DK, AB

Visualization: AP

Writing – original draft: AP, VV

Writing – review and editing: AP, SP, KS, PT, DK, PRC, AB, ABP, VV

