## Supplementary Materials for "Chronic postnatal chemogenetic activation of forebrain excitatory neurons modulates adult glial function and metabolism in male mice"

**Figure S1**

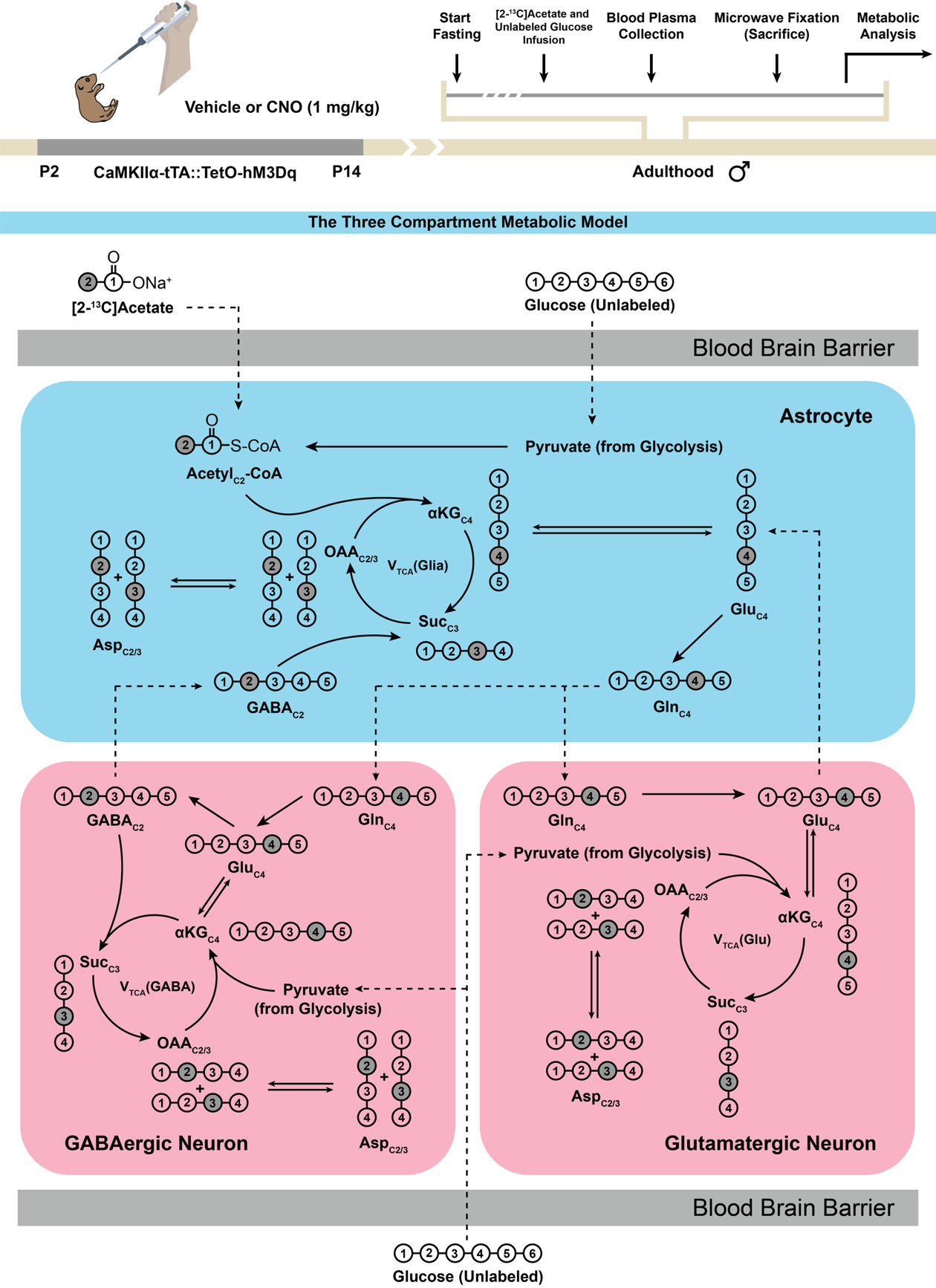

**Figure S1: A three-compartment metabolic model depicting ^13^C labeling of various metabolites in the forebrain from [2-^13^C]acetate and unlabeled glucose.** Shown is a schematic showing preferential uptake and further oxidation of [2-^13^C]acetate in astrocytes and subsequent neurotransmitter cycling between astrocytes, glutamatergic neurons, and GABAergic neurons. [2-^13^C]Acetate is preferentially metabolized in astrocytes to produce Gln_C4_ via the TCA cycle, which is further transported to glutamatergic and GABAergic neurons. Unlabeled glucose acts as the substrate for glycolysis in astrocytes and neurons. Gln_C4_ is subsequently metabolized into neuronal Glu_C4_ and GABA_C2_ via glutamine-glutamate and glutamine-GABA cycles operating between astroglia and glutamatergic neurons or astroglia and GABAergic neurons, respectively. Only the first round of TCA and neurotransmitter cycling is depicted here for simplicity. Other versions of ^13^C-labeled metabolites, such as Glu_C3_, Glu_C2_, GABA_C4_, and Gln_C2_, are produced in subsequent rounds of TCA and neurotransmitter cycling. Abbreviations: CoA – coenzyme A; α-KG – α-ketoglutarate; Suc – succinate; OAA – oxaloacetate; Asp – aspartate; Gln – glutamine; Glu – glutamate; GABA – γ-aminobutyric acid; TCA – tricarboxylic acid; V_TCA_(Glia) – astroglial TCA cycle flux; V_TCA_(GABA) – GABAergic TCA cycle flux; V_TCA_(Glu) – glutamatergic TCA cycle flux.

**Figure S2**

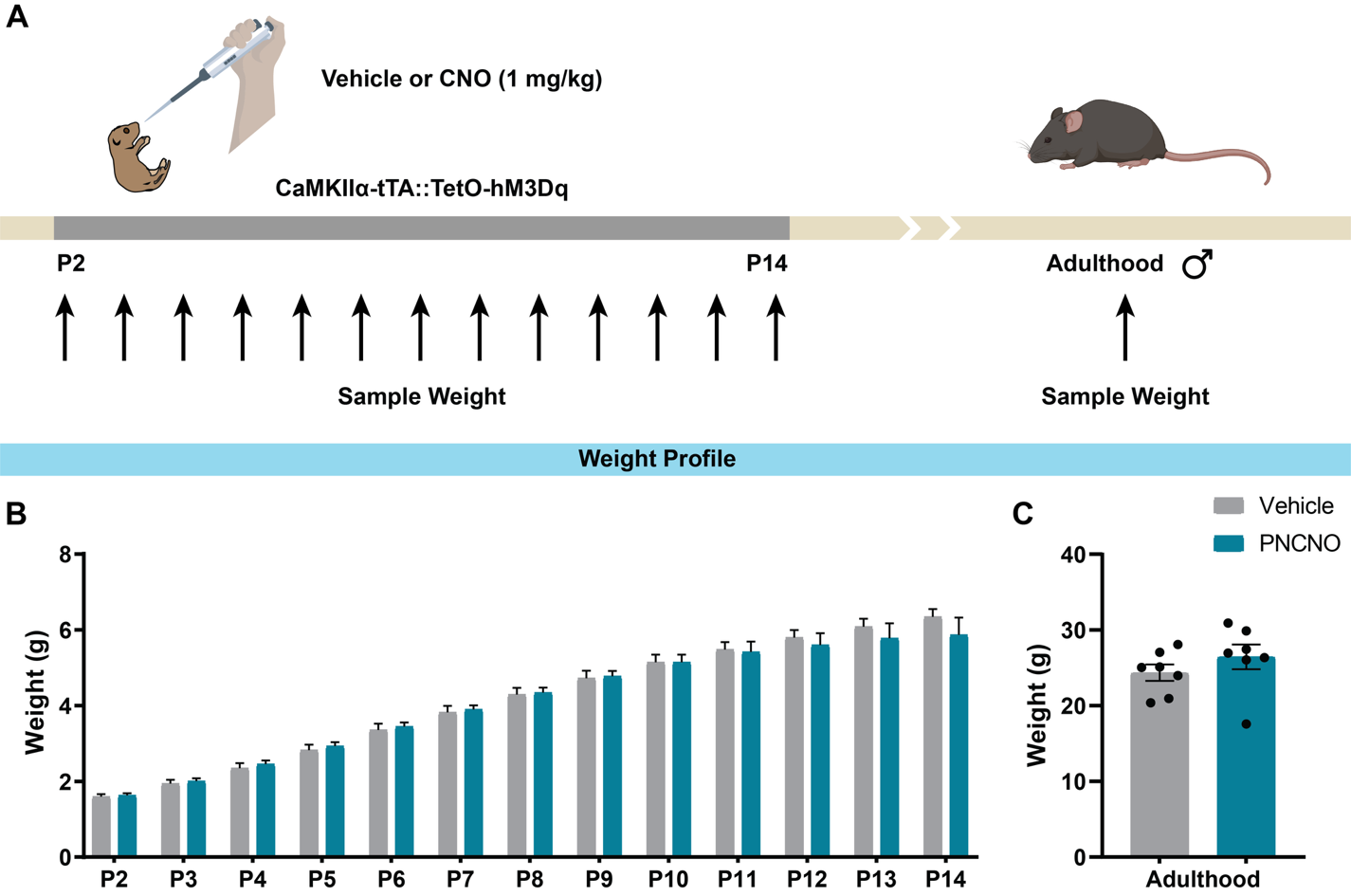

**Figure S2: Chronic CNO administration during the postnatal window does not alter the weight of bigenic CaMKIIα-tTA::tetO-hM3Dq pups during CNO administration or in adulthood. (A)** Shown is a schematic of the experimental paradigm used to determine the influence of chronic oral administration of CNO (1 mg/kg) in bigenic CaMKIIα-tTA::tetO-hM3Dq mouse pups from P2 to P14 on gross weight during the drug treatment (P2 to P14) and in adulthood. **(B)** Bigenic CaMKIIα-tTA::tetO-hM3Dq mouse pups orally administered with CNO (1 mg/kg) once daily from P2 to P14 did not differ in gross weight across the duration of drug treatment compared to the vehicle-treated controls (n = 7 litters per group; 6-8 pups per litter). **(C)** Bigenic CaMKIIα-tTA::tetO-hM3Dq adult male mice with a history of chronic PNCNO (1 mg/kg) treatment from P2 to P14 did not show any significant change in gross weight compared to vehicle-treated controls (n = 7 animals per group). Results are expressed as the mean ± S.E.M. For longitudinal body weight analysis from P2 to P14, groups are compared using two-way repeated measures ANOVA with Greenhouse-Geisser correction. For adult weights, groups are compared using the two-tailed, unpaired Student’s *t*-test. Abbreviations: PNCNO – postnatal clozapine N-oxide.

**Figure S3**

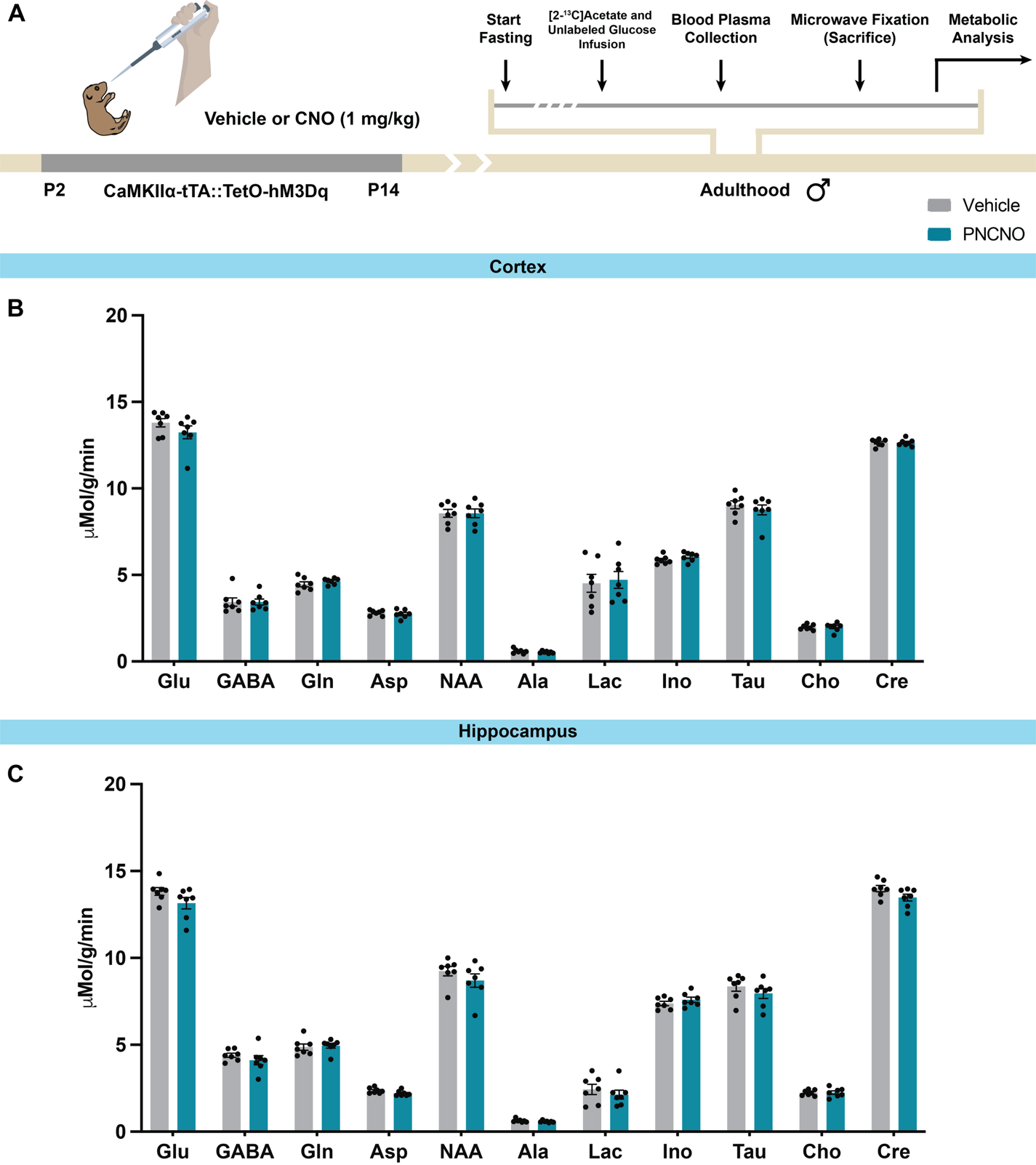

**Figure S3: Concentration of metabolites in neocortical and hippocampal tissues derived from adult bigenic CaMKIIα-tTA::tetO-hM3Dq male mice with a history of PNCNO treatment. (A)** Shown is a schematic of the experimental paradigm where bigenic CaMKIIα-tTA::tetO-hM3Dq mouse pups were orally administered vehicle or CNO (1 mg/kg) from P2 to P14 and then left undisturbed until adulthood prior to subjecting adult male mice to metabolic analysis using (^1^H-(^13^C)) NMR spectroscopy. (^1^H-(^13^C)) NMR spectra of neocortical and hippocampal tissue extracts were recorded and analyzed as previously described (Patel et al., 2001; Saba et al., 2017). **(B, C)** Shown are graphs containing the concentration of different metabolites from non-edited (^1^H-(^12^C + ^13^C)) NMR spectra measured against [2-^13^C]glycine as reference. We do not see a significant change in the concentration of these metabolites within the neocortex **(B)** and the hippocampus **(C)** of PNCNO-treated bigenic CaMKIIα-tTA::tetO-hM3Dq adult male mice, compared to their vehicle-treated controls (n = 7 per group). Results are expressed as the mean ± S.E.M., and groups are compared using the two-tailed, unpaired Student’s *t*-test. Abbreviations: PNCNO – postnatal CNO; Glu – glutamate; GABA – γ-aminobutyric acid; Gln – glutamine; Asp – aspartate; NAA – N-acetylaspartate; Ala – alanine; Lac – lactate; Ino – inositol; Tau – taurine; Cho – choline; Cre – creatine.
